# Parameter non-identifiability limits ecological inference from historical plague mortality records

**DOI:** 10.64898/2026.09.26.754648

**Authors:** Nich W. Martin, Nicolas Gauthier

## Abstract

Historical plague mortality records provide unusually detailed temporal records of epidemic dynamics. However, the ecological processes underlying those dynamics are rarely observed. This creates a fundamental challenge for inference into ecological mechanisms, as multiple combinations of unobserved ecological parameters and initial conditions may produce similar mortality trajectories. We investigated this problem by examining the geometry of parameter compensation for models of the Barcelona plague outbreak of 1490. We fit a mechanistic rat-flea-human transmission model to daily human mortality data under three values of flea search efficiency, then characterized the resulting joint posterior distributions. We then used a phenomenological model in which human infection was driven by a time-varying external force of infection, allowing us to separate parameter dependencies arising from deterministic structure. The mechanistic model reproduced the broad temporal structure of the observed epidemic, but exhibited strong linear and nonlinear dependencies among transmission parameters and initial rat and flea conditions. Similar dependencies persisted in the phenomenological model, despite its substantially reduced complexity. However, fixing the timing of the external hazard substantially reduced these dependencies. These results demonstrate that detailed mortality records can constrain the temporal pattern of an epidemic without uniquely identifying the mechanisms that generated it. Rather than simply adding additional mortality records, improving mechanistic inference from historical epidemics may require recovering independent observations of upstream ecological processes, such as timing of rodent or flea dynamics as well as environmental drivers. Mechanistic models can therefore serve not only to reconstruct historical epidemics but also to identify which missing observations would most improve inference.

## Introduction

Historical plague (*Yersinia pestis*) epidemics provide a unique opportunity to study infectious disease dynamics before the development of modern epidemiological surveillance and antibiotic use. Written records, demographic data, and mortality registers document outbreaks across centuries and geographic regions, and in some cases provide mortality observations at daily or weekly temporal resolution (Roosen and Curtis, 2018). These records have supported extensive reconstruction of the timing, magnitude, and spatial distribution of historical plague epidemics (Wood et al., 2003; Christakos and Olea, 2005). Yet the biological processes that generated those patterns of mortality remain difficult to estimate.

While detailed mechanistic epidemic models are attractive, presumably linking observed epidemic trajectories to biological processes, data available for historical outbreaks are substantially simpler than the ecological systems represented by these models (Park et al., 2018). Most historical plague models rely primarily on daily or weekly records of human mortality (e.g. Dean et al., 2018). However, these observations provide little direct information about the ecological processes occurring upstream of human infection (Park et al., 2018; Earn et al., 2020). Thus, a fundamental mismatch exists between the ecological complexity represented by the model and the information contained in the observations, as upstream, latent processes must be inferred indirectly from the patterns they produce.

This mismatch is central to the problem of parameter identifiability (Cobelli and Distefano, 1980). A parameter within a model is uniquely identifiable when available observations provide sufficient information to distinguish its value from alternative values. Parameter non-identifiability arises from equifinality, where different combinations of the same parameters generate the same or similar observable trajectories (Marschmann et al., 2019; Gallo et al., 2022). Non-identifiability is a recognized problem in infectious disease modeling (Wang, 2026), particularly for systems in which transmission depends on interactions among multiple populations. Even relatively simple vector-borne disease models can exhibit substantial parameter dependence despite observations of both hosts and vectors (Kao and Eisenberg, 2018). Historical plague presents an especially severe version of this problem. Human mortality may be recorded in considerable temporal detail, while observations of the ecological processes represented by a mechanistic model—such as rodent abundance, flea abundance, infection prevalence, or transmission among host species—are rarely available at comparable temporal resolution (Antoine, 2008). Increasing model complexity does not resolve this ambiguity. In fact, if additional biological processes are represented without corresponding observations, additional parameters may create additional compensatory pathways through which alternative mechanisms can lead to similar outbreaks in different ways. These compensatory pathways, when present, have unique, qualitative geometries when comparing one parameter or initial state against another.

Here, we investigate the geometry of these compensations using the historical number of human deaths from the 1490 Barcelona plague outbreak and analyze the data using two complementary models. First, we fit a mechanistic rat–flea–human model adapted from Keeling and Gilligan (2000) and Dean et al. (2018), examining how alternative values of flea search efficiency affects the compensatory geometry between transmission rates, initial number of infected rats, and size of the rat population at risk. Second, we construct a phenomenological model in which the entire rat–flea transmission system is represented as an exogenous, time-varying force of infection acting on humans. This model removes much of the explicit ecological structure and provides visualizations of the qualitative features of parameter dependence. Finally, we ask whether independent information about the timing of the external hazard (peak vector infection) can decrease some of these parameter dependencies.

## Methods

### Data

We analyzed daily human mortality data from the 1490 plague outbreak in Barcelona, derived from the Barcelona bills of mortality (Smith, 1936), as digitized by Dean et al. (2018) and made available through the *yersinia* R package (Gauthier and Martin, 2026). The outbreak is estimated to have occurred in a population of approximately 25,000 people (Dean et al., 2018). The mortality record provides a temporally resolved observation of the downstream human epidemic but contains no direct observations of rat, *Rattus rattus*, or flea populations represented in the mechanistic model.

### Mechanistic Model

Our Rat-Flea-Human transmission model was adapted from Keeling and Gilligan (2000) and Dean et al. (2018), specifically ignoring features such as natural rat and human growth and death rates, focusing purely on transmission and mortality parameters. The model uses three population compartments: rat, flea, and human; where our rat compartment is represented by:

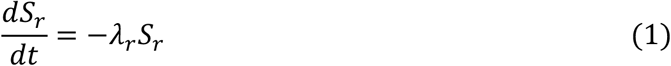

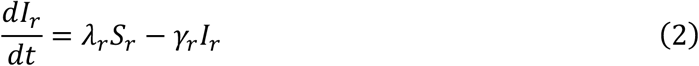

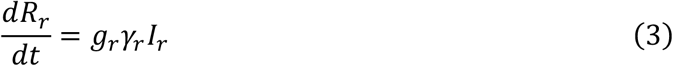

and

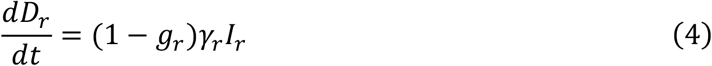

where *S*_*r*_ represent the population of susceptible rats, *I*_*r*_ represents infected rats, *R*_*r*_ represents recovered rats, *D*_*r*_ represents dead rats, *λ*_*r*_ represents the density-dependent flea-to-rat transmission rate (derived below), and *γ*_*r*_ represents the death rate among infected individuals assuming a 90% probability of mortality.

Our flea compartment is represented by:

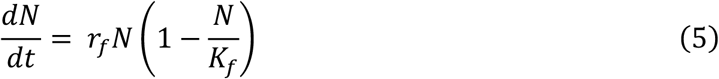

and

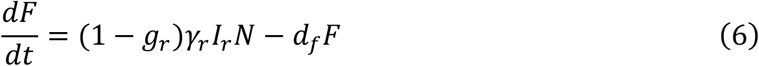

where *N* represents the average number of fleas per rat, and *F* represents the number of free fleas in the environment which emerge after an infected rat dies.

The parameters *r*_*f*_ and *d*_*f*_ represent hosted flea birth and free flea death rates, respectively, and *K*_*f*_ represents the carrying capacity of fleas per rat. The flea-rat transmission state, *λ*_*r*_, was generated from *F* such that,

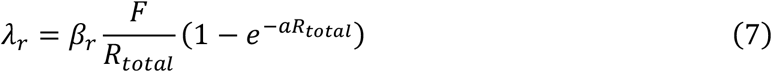

where *β*_*r*_ is the base flea-to-rat transmission rate, *R*_*total*_ is the total number of rats, where,

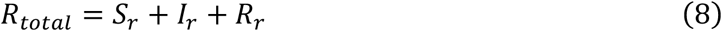

and *a* is the flea search efficiency parameter, which determines how effective free fleas are at finding a rat host. Here we treat *a* similar to Dean et al. (2018) where,

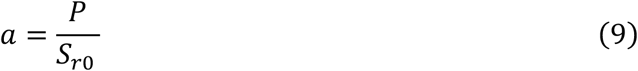

and

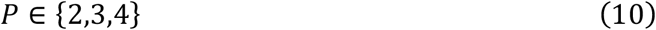

with *P* = 2, *P* = 3, and *P* = 4 representing low, medium, and high relative values, respectively. Our human compartment is as follows:

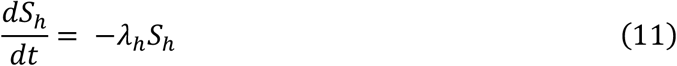

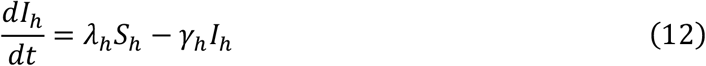

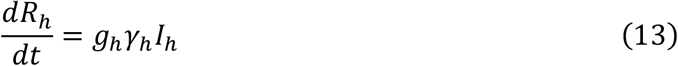

and

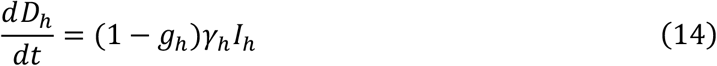

where

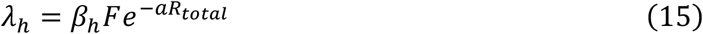

Again, *β* and *γ* represent baseline transmission and death among infected rates, respectively. A number of our parameter values were fixed for model simplicity using the same values as Dean et al. (2018) and can be found in table 1.

**Table 1.**
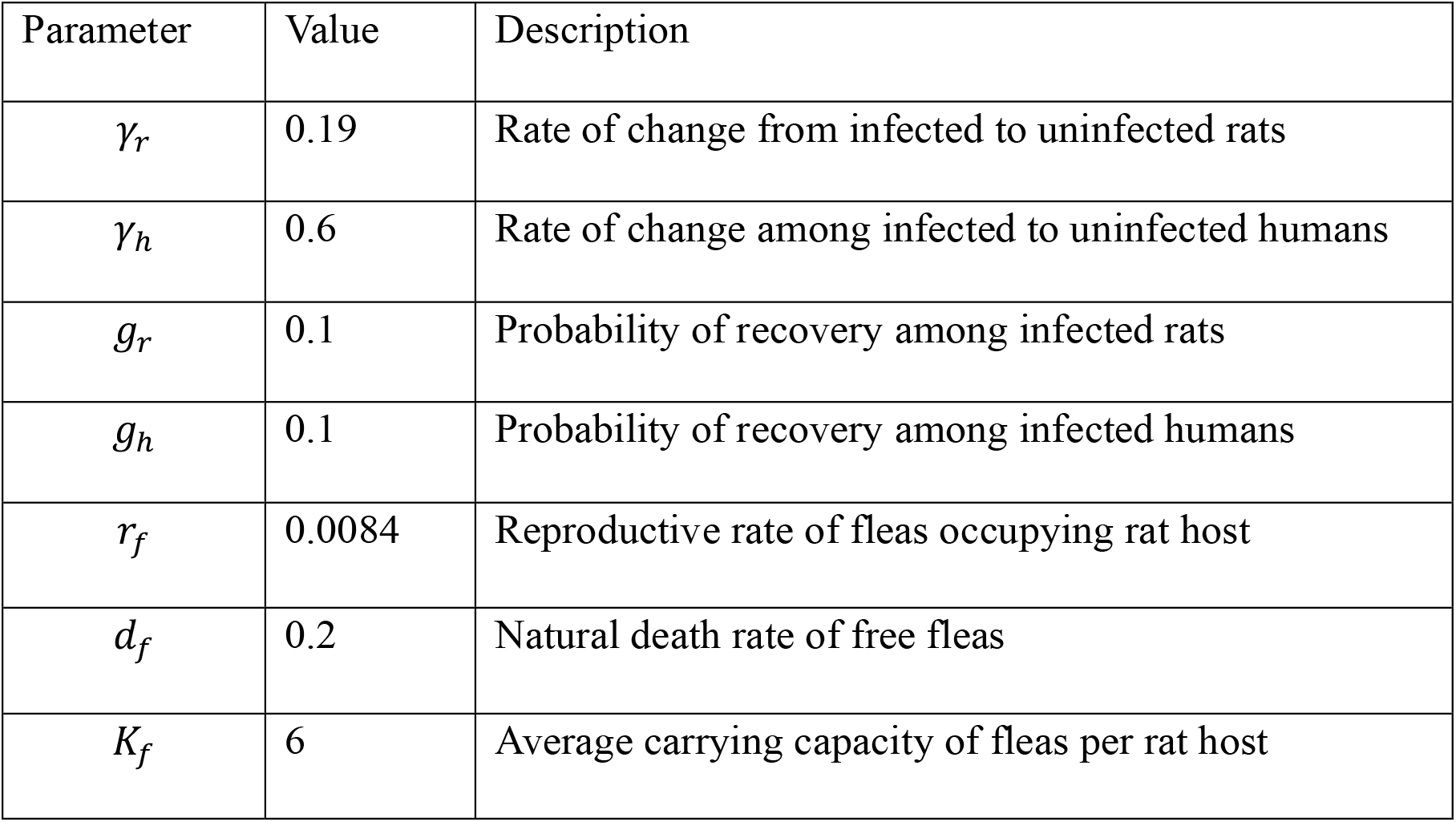
Fixed parameters of the rat-flea-human model.

We fit the model to daily human mortality using a negative-binomial likelihood to accommodate for overdispersion as suggested by Park et al. (2018). We used Hamiltonian Monte Carlo implemented in Stan (2.37) to estimate *β*_*r*_, *β*_*h*_, the initial number of infected rats, 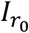, the population of rats at risk 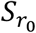, as well as the error term of negative-binomial likelihood function. The primary objective of this analysis was not to obtain point estimates of the ecological parameters, but to obtain their joint posterior geometry.

### Exogenous force-of-infection model

Our exogenous, time-varying force of infection model was designed to represent the downstream effect of a rat–flea transmission system without explicitly modeling rats or fleas, where force of infection *λ*(*t*) representing the per-susceptible, per-day risk of infection arising from the rat–flea system. Here *λ*(*t*) is parameterized as a Gaussian burst (or pulse) with total area *A*, peak time *t*_*peak*_, and width *σ*, such that,

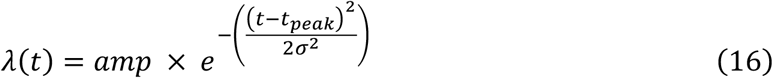

and the instantaneous peak height (amplitude) is derived as,

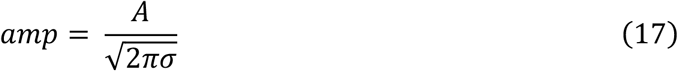

Biologically, *A* is the cumulative spillover pressure over the outbreak, integrating infectious fleas, bite rates, and transmission rate; *t*_*peak*_ marks when spillover pressure is highest (e.g., near the peak of the rat epidemic); and *σ* controls duration of the hazard, reflecting rat epidemic length, flea survival off-host, season/climate, and human behavior.

Human infection dynamics follow a simple deterministic system with states *S*(*t*) (susceptible), *I*(*t*) (infected), and *C*_*d*_ (cumulative deaths). The ordinary differential equations are:

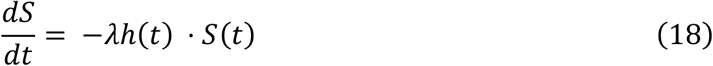

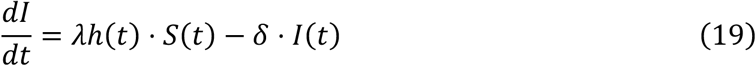

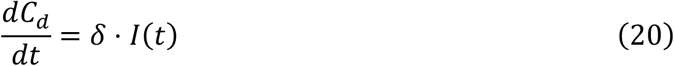

where δ is the combined effect of infection duration and probability of an infected individual dying. Again, likelihood for daily deaths were modeled using a negative-binomial distribution to accommodate overdispersion.

This phenomenological model allowed us to separate two questions that are confounded in the mechanistic formulation. First, can mortality data alone constrain the magnitude, duration, and timing of an effective external hazard? That is, can these parameters be uniquely identified for an outbreak for the purposes of inference? Second, how does this geometry change when independent information about hazard timing is available? We fit two versions of the phenomenological model. One where *t*_*peak*_ is estimated, and one where it is fixed at the mean estimated value from the first model, an experiment designed to isolate whether dependencies among the remaining parameters are intrinsic or induced by uncertainty in hazard timing. This approach is motivated by the broader observation that identifiability can depend strongly on which components of a dynamical system are observed and how completely the system trajectory is observed (Sauer et al., 2022; Chowell et al., 2023). Priors and initial conditions for all model parameters can be found in the supplementary materials.

## Results

The rat-flea-human model reproduced the general structure of the observed Barcelona epidemic (S1). However, joint posterior distributions revealed a combination of linear, nonlinear, and no apparent dependencies among estimated parameters (Fig. 1). In particular, flea-to-rat transmission rate, *β*_*r*_, showed little to no association with the other estimated parameters and occupied distinct, non-overlapping regions of the parameter space for different values of *P* (low: 0.066-0.072; medium: 0.056-0.060; and high: 0.050-0.054), increasing with higher values of *P*.

**Figure 1.**
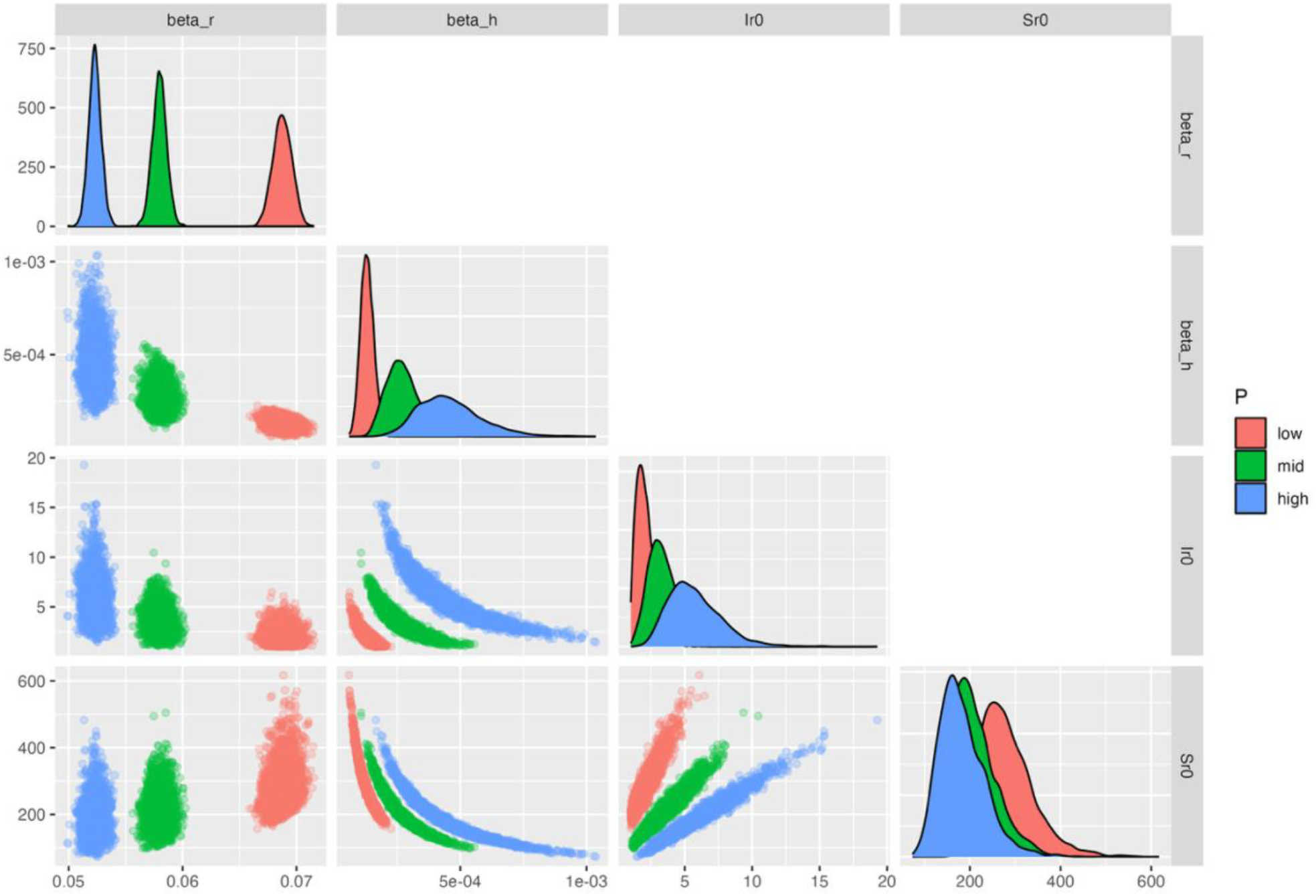
Joint posterior distributions of the rat-flea-human model for flea-to-rat transmission (beta_r), flea-to-human transmission (beta_h), initial number of infected rats (Ir0), and population of rats at risk (Sr0).

In contrast, flea-to-human transmission, *β*_*h*_, showed noticeably non-linear dependencies with the initial number of infected rats, 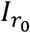, and at-risk population of rats, 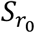, where both initial states decreased with increasing values of *β*_*h*_. This non-linear geometry is consistent with multiplicative parameter compensation, where the trajectory is observed given *ψ*_*X*_ and *ψ*_*X*_ = *X* ⋅ *β*_*h*_ where 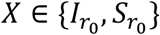. Both 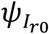 and 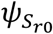 took on a unique range of values, increasing with increasing values of *P* (Table 2). Unlike their relationship with *β*_*h*_ the dependency between 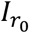 and 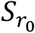 was distinctly linear, suggesting an additive compensation, with the slope appearing to decrease with increasing values of *P* (low = 77.32, medium = 40.94, high = 25.26).

**Table 2.** Ranges of joint posterior combinations for different values of *P*, where low: *P* = 2, medium: *P* = 3, and high: *P* = 4.

| Parameter combination | Low | Medium | High |
| --- | --- | --- | --- |
| $I_{r_0} \cdot \beta_h$ | $(1.40:4.11) \cdot 10^{-4}$ | $(4.84:12.7) \cdot 10^{-4}$ | $(15.0:33.4) \cdot 10^{-4}$ |
| $S_{r_0} \cdot \beta_h$ | $(3.15:3.97) \cdot 10^{-2}$ | $(4.68:6.02) \cdot 10^{-2}$ | $(6.91:8.79) \cdot 10^{-2}$ |
| $S_{r_0} - I_{r_0}$ | 156:611 | 96.9:495 | 71.7:463 |

Our exogenous hazard model reproduced both the broad general and temporal structure of the mortality data (S2), using substantially fewer parameters, and without representing rats and fleas explicitly. Nevertheless, the joint posterior distributions continued to exhibit parameter dependence (Fig. 2). In fact, only cumulative spillover pressure, *A*, did not show noticeable dependencies with the other parameters, while peak spillover pressure, *t*_*peak*_, duration hazard, *σ*, and the combined duration of infection and probability of mortality, *δ*, all showed positive relationships. The relationship between *t*_*peak*_ and *σ* was noticeably linear, while the relationships between those parameters and *δ* formed a characteristic ridge. However, when we assumed a known value for *t*_*peak*_ using our point prior, the relationship between the *σ* and *δ* joint posteriors appeared less structured as well as less clustered (Fig. 3).

**Figure 2.**
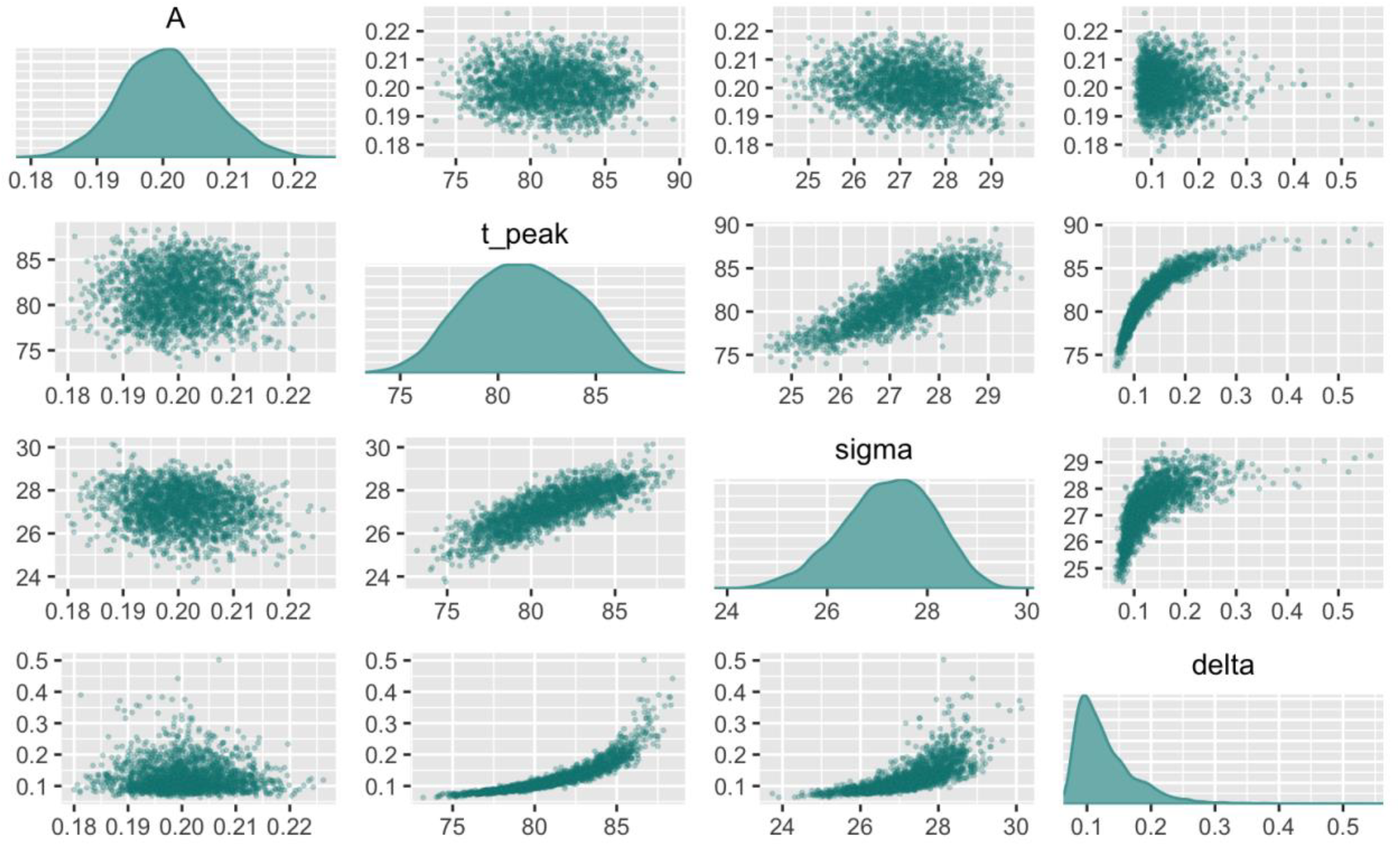
Joint posteriors of the phenomenological model for the cumulative spillover outbreak pressure (*A*), peak spillover pressure (*t*_*peak*_), duration hazard (*σ*), and mortality rate of infected (*δ*).

**Figure 3.**
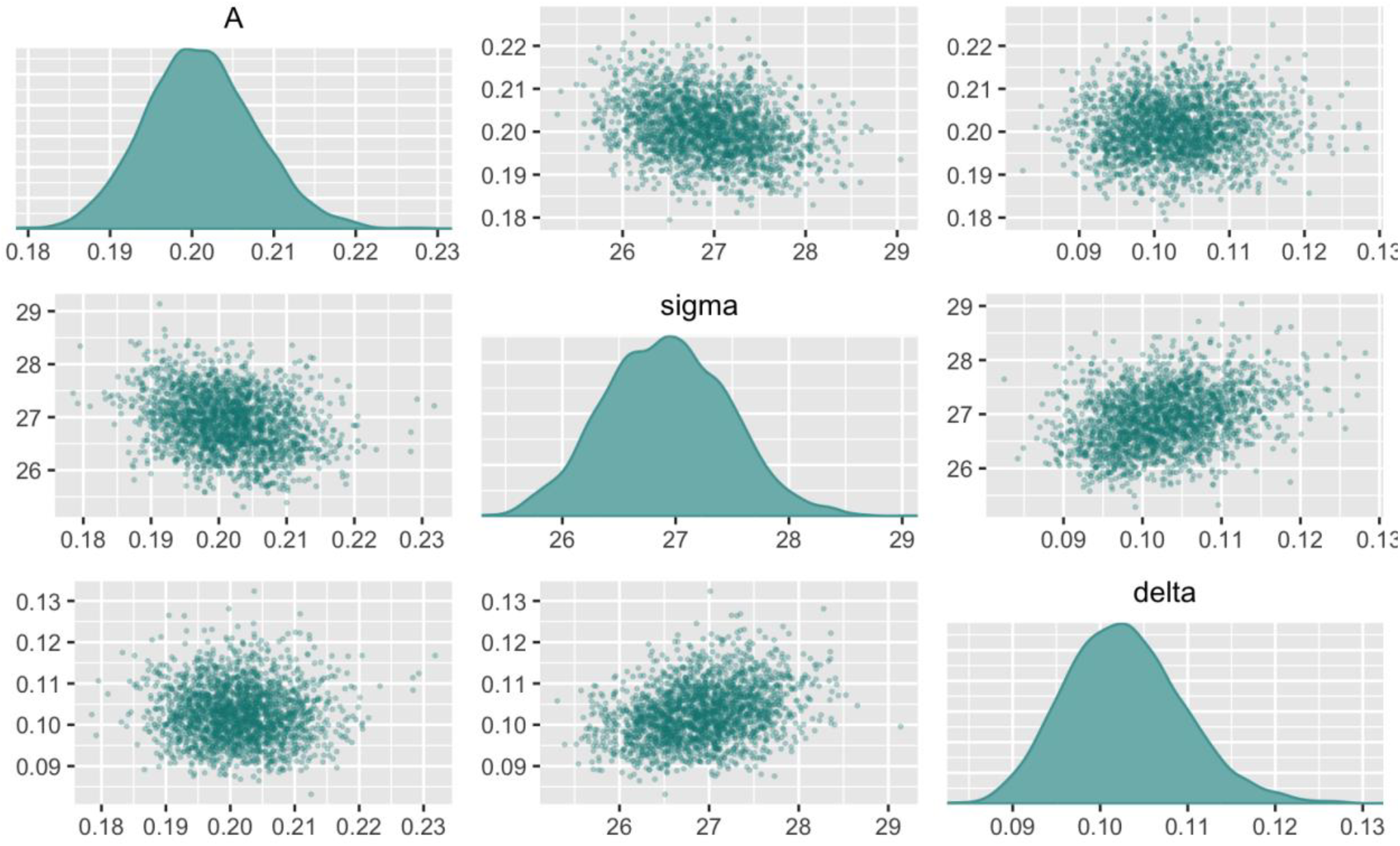
Joint posteriors of the phenomenological model for the cumulative spillover outbreak pressure (*A*), duration hazard (*σ*), and mortality rate of infected (*δ*) with peak spillover pressure (*t*_*peak*_) fixed at 80.

## Discussion

Historical plague mortality records can provide remarkably detailed information about the temporal trajectory of an epidemic. That information does not necessarily extend to the ecological processes that produced the trajectory. The rat-flea-human model reproduced the observed Barcelona mortality epidemic while permitting substantial variation in transmission rates and initial rat conditions. Varying combinations of these parameters occupied narrow, linear and nonlinear regions of posterior space, such that mortality data could not distinguish among multiple configurations of the underlying transmission system, showing strong equifinality in the sense of Marschmann et al. (2019). Estimated flea-to-human transmission rates, *β*_*h*_, show strong associations with the posterior estimates of both the initial number of infected rats, 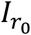 and the number of susceptible rats at risk, 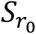, suggesting this model cannot separate whether the observed data support a relatively large initial population of infected rats with low flea-to-human transmission versus a smaller initial population of infected rats with high flea-to-human transmission. Likewise, the relationship between the posterior estimates for 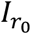 and 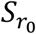 show that as the initial number of infected rats increase the number of rats at risk increases as well given the observed data, with the effect decreasing as flea search efficiency increases.

These results are consistent with a broader literature showing that parameter estimation in mechanistic epidemic models can be limited by structural and practical non-identifiability and dynamical compensation (Chowell, 2017; Sauer et al., 2022; Chowell et al., 2023). Central to our analysis is that parameter dependence persists after replacing the rat-flea system with a simple phenomenological hazard, and that those dependencies change qualitatively with additional information. There are two implications for how historical plague epidemics are modelled and what historical research might usefully target.

First, human mortality data alone constrain mechanistic plague models weakly. But historical documents may contain other observations bearing on the timing of ecological events or shifts in animal behavior. Descriptions of changes in rodent abundance, sudden disappearance of rats, unusual animal mortality, changes in flea abundance, or environmental conditions associated with the onset and decline of an outbreak could constrain components of the transmission process that mortality cannot identify alone. Historical observations need not provide complete ecological time series to be useful; observations that independently constrain specific features of the transmission process may eliminate otherwise plausible regions of parameter space. Even with such constraints, however, identifiability remains difficult. This extends Park et al.’s (2018) observation that Dean et al. (2018) treat several upstream quantities as known. Here, we show that differences in upstream point priors not only affect downstream parameter estimates, but also the geometry of the joint posterior (Fig. 1). Furthermore, model selection criteria (e.g., BIC) identify which model predicts the data most efficiently, but do not provide evidence that one causal model is more likely than another (Mac Nally, 2000; Burnham and Anderson, 2004).

Second, at the phenomenological level, our fixed *t*_*peak*_ analysis shows how information about the timing of an upstream hazard changes the geometry of the inference problem, altering dependencies among parameters that remain strongly coupled when hazard timing is inferred from mortality alone. Continuous covariates such as reconstructed climate may therefore be more informative about the drivers of plague outbreaks than additional mortality series. Lewnard and Townsend (2016) illustrate the potential of such an approach, combining archival observations of rat resistance and flea abundance with climatic drivers to explain shifts in the seasonality of outbreaks. Where comparable records can be assembled, incorporating them is likely to do more for inference than fitting additional outbreaks with the same mortality-only data. More mortality curves add replication but do not allow us to observe the rat and flea dynamics that generated them.

## Conclusions

Mortality records during plague outbreaks can tell us when an outbreak occurred and describe its features (e.g., total number of deaths, duration, etc.) without telling us why. Our results show that multiple configurations of a mechanistic rat-flea-human system can produce the same mortality trajectory, and that substantial parameter dependence persists even when that ecological system is replaced by a simple phenomenological hazard. The reduction of this dependence when hazard timing is independently specified demonstrates that additional information about upstream processes can fundamentally change what can be inferred from mortality records.

The implication is not that historical plague is inaccessible to mechanistic inference, but that historical records need to contain or be supplemented with information about mechanisms other than those we seek to infer. Mechanistic models can help identify where that information is missing and, importantly, what kinds of observations would be most valuable to recover. In this way, models of historical epidemics can do more than reconstruct past mortality: they can help guide the search for the ecological evidence needed to understand the mechanisms behind it.

## Notes

### Competing Interest Statement

The authors have declared no competing interest.

### Funding

- National Science Foundation: 2409068

