## Supplementary tables and figures for "Parameter non-identifiability limits ecological inference from historical plague mortality records"

*Rat-Flea-Human Model*

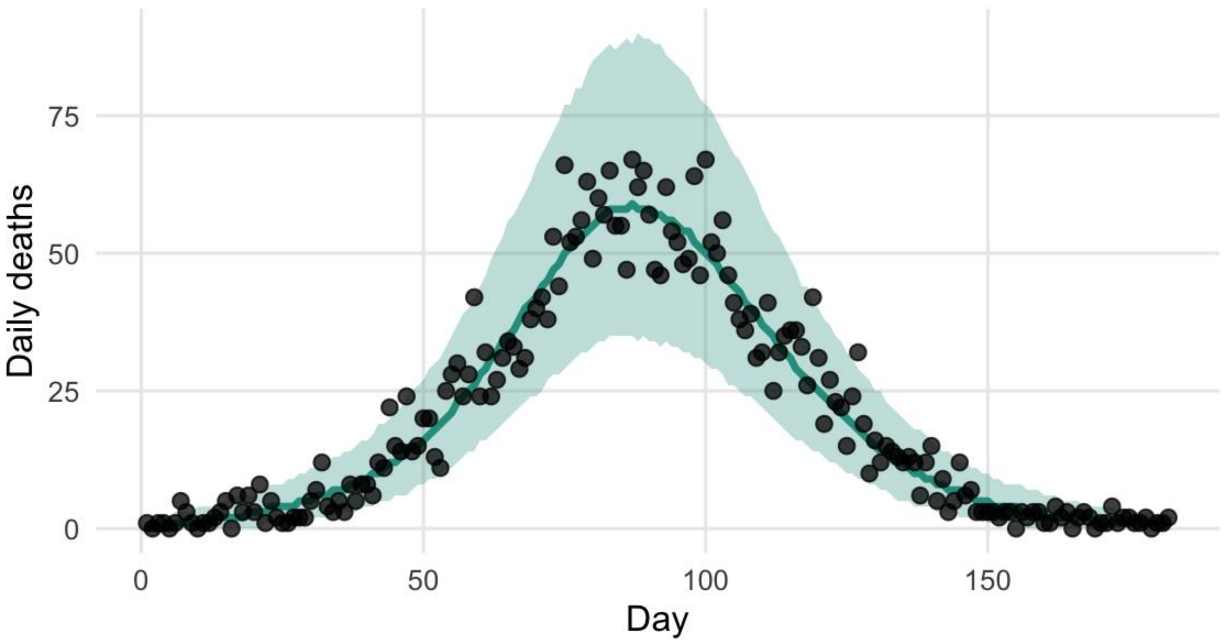


S1. Posterior predictive simulation for the number of deaths against day with observed deaths (CI 95%).

| |
|---|
| $\beta_r \sim lognormal(-3, 0.5)$ |
| $\beta_h \sim lognormal(-5, 0.5)$ |
| $I_{r_0} \sim lognormal(\, ln(5), 0.5)$ |
| $N_0 \sim lognormal(ln(5), 0.15)_{[0,7]}$ |
| $\phi \sim exponential(1)$ |
| $S_{r_0} = 5{,}000 - I_{r_0}$ |
| $F_0 = 1$ |

Supplementary Table 1. (ST1) Priors and initial states for the flea-rat-human model

```
Inference for Stan model: anon_model.
4 chains, each with iter=2000; warmup=1000; thin=1;
post-warmup draws per chain=1000, total post-warmup draws=4000.

          mean se_mean    sd    0%     50%   100% n_eff Rhat
beta_r    0.06    0.00  0.00  0.06    0.06   0.06  2043    1
beta_h    0.00    0.00  0.00  0.00    0.00   0.00  1261    1
Ir0       3.46    0.03  1.18  1.07    3.28  10.44  1398    1
S       207.12    1.36 50.26 98.18 200.04 504.75  1368    1

Samples were drawn using NUTS(diag_e) at Mon Sep 21 12:55:42 2026.
For each parameter, n_eff is a crude measure of effective sample size,
and Rhat is the potential scale reduction factor on split chains (at
convergence, Rhat=1).
```

ST2. Summary of posteriors and convergence metrics $a = mid$

*Exogenous Model*

| |
|---|
| $A \sim lognormal(ln(0.3), 0.5)$ |
| $t_{peak} \sim uniform(0, t_{max})$ |
| $\sigma \sim lognormal(ln(5), 0.5)$ |
| $\delta \sim lognormal(ln(0.10), 0.4)$ |
| $\phi \sim exponential(1)$ |

ST3. Exogenous model priors

```
Inference for Stan model: anon_model.
4 chains, each with iter=2000; warmup=1000; thin=1;
post-warmup draws per chain=1000, total post-warmup draws=4000.

        mean se_mean   sd    5%   50%   95% n_eff Rhat
A       0.18    0.00 0.01  0.17  0.18  0.19  2356    1
t_peak 81.15    0.08 2.73 76.74 81.10 85.75  1175    1
sigma  27.21    0.02 0.88 25.68 27.26 28.58  1292    1
delta   0.13    0.00 0.05  0.08  0.12  0.24  1220    1

Samples were drawn using NUTS(diag_e) at Tue Sep 15 10:38:34 2026.
For each parameter, n_eff is a crude measure of effective sample size,
and Rhat is the potential scale reduction factor on split chains (at
convergence, Rhat=1).
```

ST4. Summary of posteriors and convergence metrics

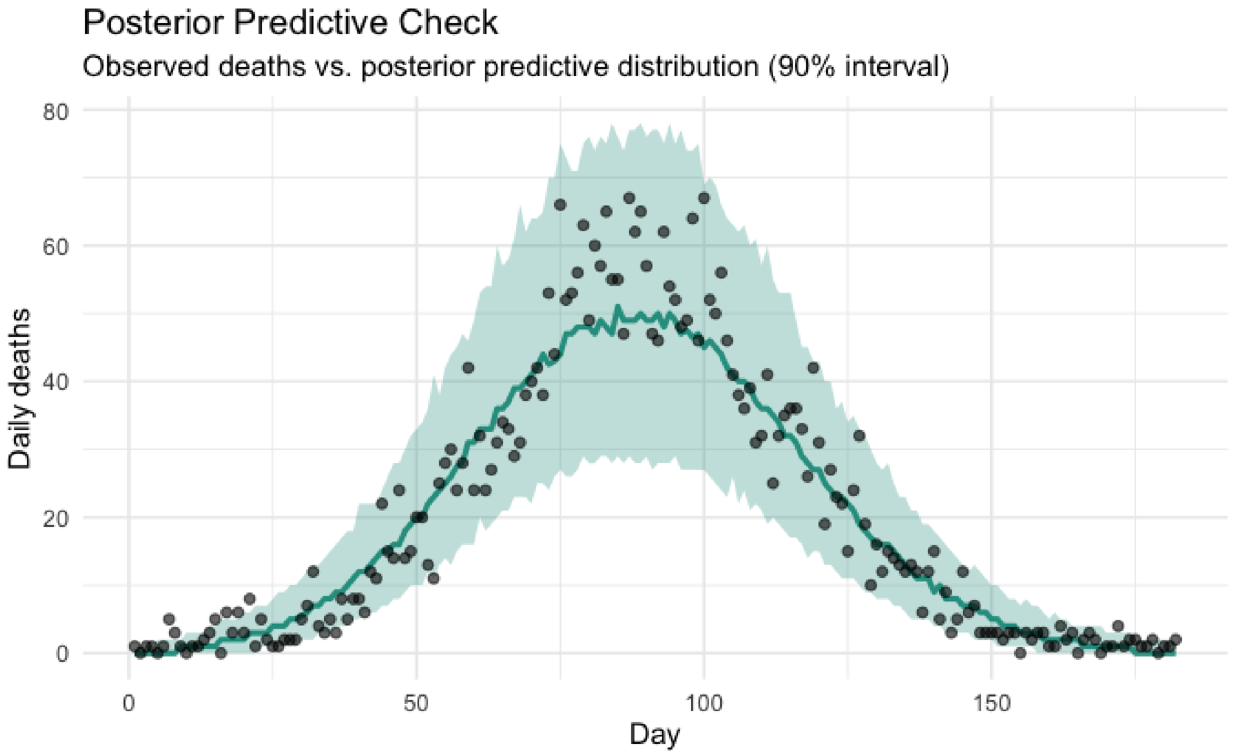


S2. Posterior predictive simulation of the exogenous model, number of human deaths per day (CI. 95%).

```
Inference for Stan model: anon_model.
4 chains, each with iter=1000; warmup=500; thin=1;
post-warmup draws per chain=500, total post-warmup draws=2000.

       mean se_mean   sd    5%   50%   95% n_eff Rhat
A      0.20    0.00 0.01  0.19  0.20  0.21  1977    1
sigma 26.88    0.01 0.53 26.03 26.87 27.74  1869    1
delta  0.10    0.00 0.01  0.09  0.10  0.11  1792    1

Samples were drawn using NUTS(diag_e) at Tue Sep 15 10:13:13 2026.
For each parameter, n_eff is a crude measure of effective sample size,
and Rhat is the potential scale reduction factor on split chains (at
convergence, Rhat=1).
```

ST5. Summary of posteriors and convergence metrics for exogenous fixed t_peak model.
